# Expanding the *Caenorhabditis elegans* auxin-inducible degron system toolkit with internal expression and degradation controls and improved modular constructs for CRISPR/Cas9-mediated genome editing

**DOI:** 10.1101/2020.05.12.090217

**Authors:** Guinevere Ashley, Tam Duong, Max T. Levenson, Michael A. Q. Martinez, Jonathan D. Hibshman, Hannah N. Saeger, Ryan Doonan, Nicholas J. Palmisano, Raquel Martinez-Mendez, Brittany Davidson, Wan Zhang, James Matthew Ragle, Taylor N. Medwig-Kinney, Sydney S. Sirota, Bob Goldstein, David Q. Matus, Daniel J. Dickinson, David J. Reiner, Jordan D. Ward

**Affiliations:** Department of Molecular, Cell, and Developmental Biology, University of California-Santa Cruz, Santa Cruz, CA 95064, USA; Institute of Biosciences and Technology, Texas A&M Health Science Center, Houston, TX 77030, USA; Department of Biochemistry and Cell Biology, Stony Brook University, Stony Brook, NY 11794, USA; Department of Biology, University of North Carolina at Chapel Hill, Chapel Hill, North Carolina, USA; Department of Molecular Biosciences, University of Texas at Austin, Austin, TX 78712, USA; Lineberger Comprehensive Cancer Center, University of North Carolina at Chapel Hill, Chapel Hill, North Carolina, USA

**Keywords:** *C. elegans*, AID system, SapTrap, self-excising cassette, CRISPR/Cas9, Transport Inhibitor Response 1

## Abstract

The auxin-inducible degron (AID) system has emerged as a powerful tool to conditionally deplete proteins in a range of organisms and cell-types. Here, we describe a toolkit to augment the use of the AID system in *Caenorhabditis elegans*. We have generated a set of single-copy, tissue-specific (germline, intestine, neuron, muscle, hypodermis, seam cell, anchor cell) and pan-somatic *TIR1-*expressing strains carrying an equimolar co-expressed blue fluorescent reporter to enable use of both red and green channels in experiments. We have also constructed a set of plasmids to generate fluorescent protein::AID fusions through CRISPR/Cas9-mediated genome editing. These templates can be produced through frequently used cloning systems (Gibson assembly or SapTrap) or through ribonucleoprotein complex-mediated insertion of PCR-derived, linear repair templates. We have generated a set of sgRNA plasmids carrying modifications shown to boost editing efficiency, targeting standardized transgene insertion sites on chromosomes I and II. Together these reagents should complement existing *TIR1* strains and facilitate rapid and high-throughput fluorescent protein::AID* tagging of factors of interest. This battery of new TIR1-expressing strains and modular, efficient cloning vectors serves as a platform for facile assembly of CRISPR/Cas9 repair templates for conditional protein depletion.

## Introduction

Conditional degrons have emerged as a powerful tool to rapidly destroy proteins of interest in order to interrogate their function in cells and in multicellular animals (Natsume and Kanemaki 2017). One of these tools, the auxin-inducible degron (AID) system, has been successfully employed in *C. elegans* (Zhang *et al*. 2015), budding yeast (Nishimura *et al*. 2009), *Drosophila* (Trost *et al*. 2016; Chen *et al*. 2018), zebrafish (Daniel *et al*. 2018), *Toxoplasma gondii* (Brown *et al*. 2017), cultured mammalian cells (Nishimura *et al*. 2009; Holland *et al*. 2012; Natsume *et al*. 2016), and mouse oocytes (Camlin and Evans 2019). The system is comprised of two components. First, a plant F-box protein named Transport Inhibitor Response 1 (TIR1) is expressed under the control of a promoter with a defined expression pattern (Figure 1). TIR1 can then interact with endogenous Skp1 and Cul1 proteins to form a functional SCF E3 ubiquitin ligase complex (Figure 1). Second, an auxin-inducible degron (AID) sequence from the IAA17 protein is fused to a protein of interest (Figure 1) (Nishimura *et al*. 2009; Natsume and Kanemaki 2017). While the full length IAA17 sequence is 229 amino acids, minimal AID tags of 44 amino acids (AID*) and 68 amino acids (mAID) have been developed (Morawska and Ulrich 2013; Li *et al*. 2019). Addition of the plant hormone, auxin, promotes TIR1 binding to the degron, leading to the ubiquitination and subsequent proteasome-mediated degradation of the degron-tagged protein (Figure 1). In *C. elegans*, the *Arabidopsis thaliana* TIR1, AID*, and mAID sequences are used (Zhang *et al*. 2015; Negishi *et al*. 2019), as this plant grows at a temperature range more similar to *C. elegans*; rice (*Oryza sativa*)-derived sequences are used in other systems (Nishimura *et al*. 2009; Natsume *et al*. 2016; Natsume and Kanemaki 2017).

**Figure 1.**
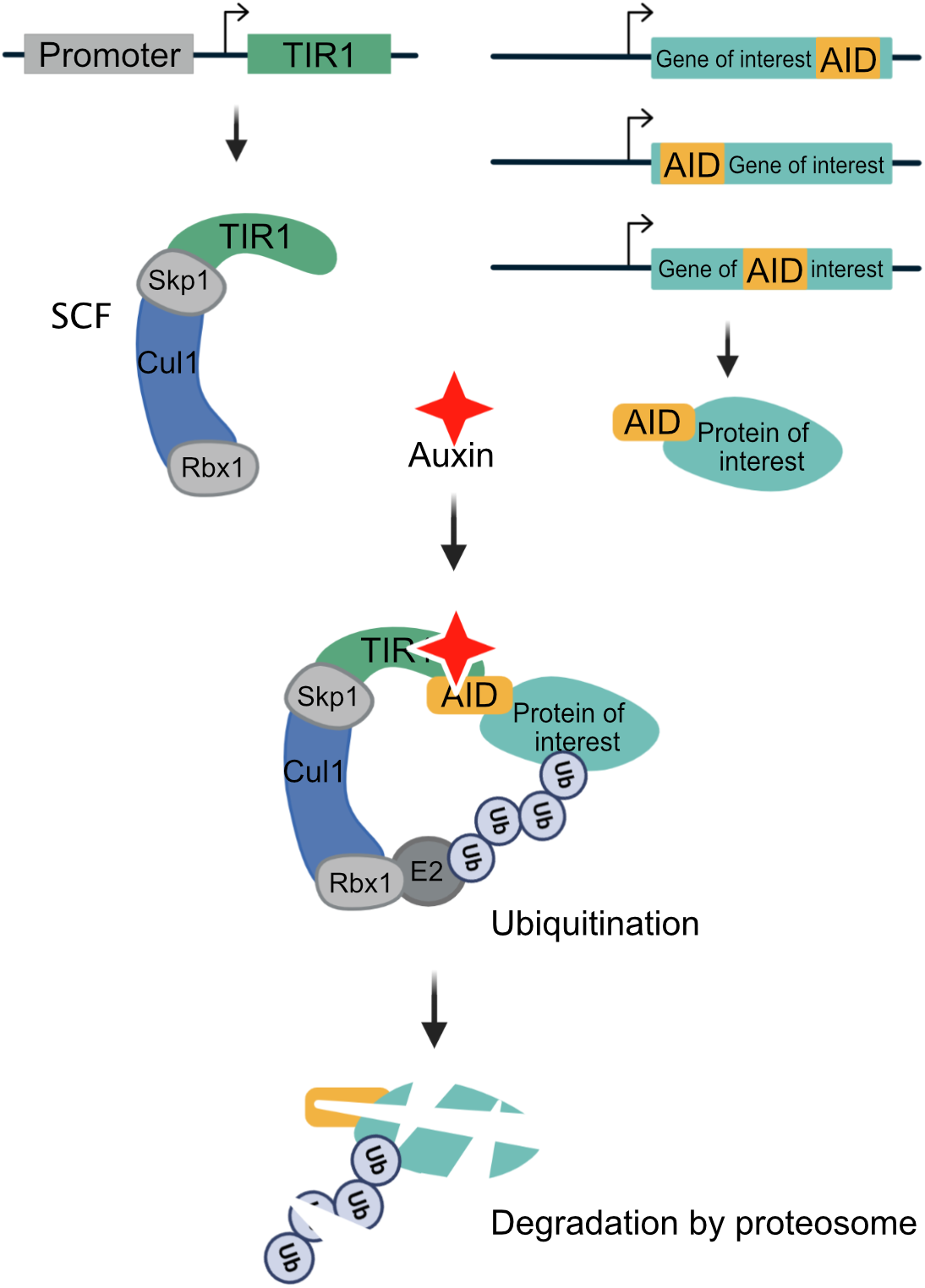
Schematic of the auxin-inducible degron (AID) system. The plant F-box protein TIR1 is expressed using a promoter of interest with a desired spatiotemporal expression pattern. TIR1 interacts with endogenous Skp1 and Cul1 proteins to form an SCF E3 ubiquitin ligase complex. An auxin-inducible degron sequence (AID) is fused to a protein of interest. We use a minimal, 44 amino acid degron sequence (AID*), but a full-length 229 amino acid AID tag or a 68 amino acid mini AID (mAID) are used in other systems. In the presence of the plant hormone auxin, TIR1 recognizes and binds the AID sequence, leading to ubiquitination and subsequent degradation of the AID-tagged protein. In *C. elegans*, the system is frequently used with single-copy TIR1 transgenes inserted into neutral loci, and AID* knock-ins into genes of interest, though extrachromosomal arrays can also be used.

**Figure 2.**
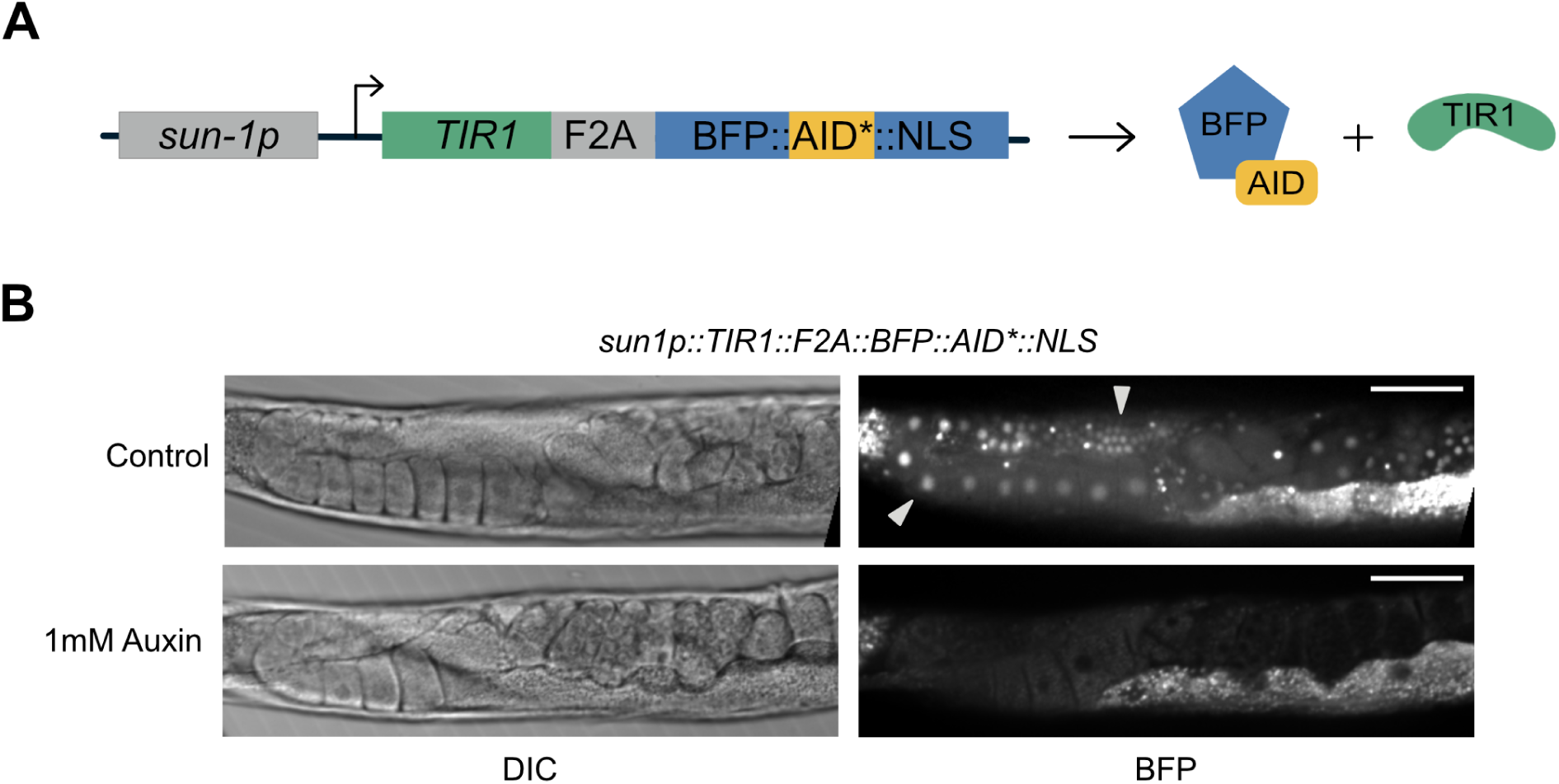
A new TIR1 expression system allows assessment of TIR1 expression and activity. A) The new TIR1 expression construct contains a *TIR1::F2A::BFP::AID*::cMyc-NLS* transgene cassette. An F2A skip sequence results in expression of two separate protein products: 1) TIR1 which will interact with endogenous SCF proteins to produce an E3 ubiquitin ligase complex, which can only bind the AID sequence in the presence of auxin; and 2) an AID*-tagged BFP protein with a cMyc nuclear localization signal (NLS) that functions as a readout for TIR1 expression and activity. The use of BFP as a reporter makes this construct compatible with simultaneous GFP and RFP imaging. B) Adult animals expressing *sun1p::TIR1::F2A::BFP::AID*::NLS*. A control animal expresses AID*-tagged BFP in the nuclei of germline and embryonic cells (white arrows). When animals are exposed to 1 mM auxin, BFP expression is undetectable. BFP channel and DIC images are provided for each condition. Note that the fluorescence signal at the lower right-hand side of each BFP image is due to intestinal autofluorescence. Scale bars represent 50 µm.

Here, we describe a new set of strains and reagents for *C. elegans* that complement the tools originally described by Zhang *et al*. (2015). We have generated a set of strains that express single-copy, tissue-specific, all-in-one TIR1 cofactor, blue fluorescent *TIR1::F2A::mTagBFP2::AID*::NLS* transgenes to enable protein depletion combined with imaging. The mTagBFP2 signal (hereafter referred to simply as “BFP”) serves as a built-in reporter and internal control for both TIR1 expression and AID-dependent activity. For many of the tissue-specific promoters driving this construct, we have created strains with insertions into standardized, neutral target sites on both chromosomes I and II (Frøkjaer-Jensen *et al*. 2008; Frøkjær-Jensen *et al*. 2012), to facilitate crossing schemes. These strains expand experimental possibilities with green and red fluorescent proteins (FPs) of interest. For example, one could deplete a green FP::AID*-tagged protein and test the effect on a red FP-tagged protein (or vice versa), or simultaneously deplete green FP::AID*- and red FP::AID*-tagged proteins of interest. One could also test the depletion of a non-tagged factor (e.g., via RNAi or CRISPR/Cas9-based mutagenesis) and monitor the levels and subcellular localization of a green or red FP::AID*-tagged proteins prior to auxin addition. We have also generated constructs to introduce FP::AID* tags into genes of interest using conventional genome editing approaches. We describe plasmid-based constructs, self-excising cassette selection vectors generated with either Gibson cloning or SapTrap, and injection of linear repair templates and Cas9 ribonucleoprotein complexes.

## Methods

### Molecular Biology

Unless otherwise stated, PCR was performed with Phusion polymerase purified in-house and 5x Phusion Green Buffer (Thermo Fisher, F538L). In the below constructs, flexible linker sequences ranging from 5-9 glycine/serine residues are used to separate cassettes within constructs, i.e. AID*, 3xFLAG, 3xMyc, TEV protease recognition sites, etc. Unless otherwise specified, pJW plasmids (Ward lab) were generated by Gibson cloning using an in-house made master mix, as described (Gibson *et al*. 2009). For two-fragment Gibson cloning 0.63 µl of each DNA fragment was mixed with 3.75 µl of the Gibson master mix and incubated for 1-4 hours at 50ºC. Longer reaction times were used for inefficient assemblies. Reactions were then transformed as described in the supplemental methods or stored at −20ºC. Detailed methods describing construct generation are provided in supplemental methods (Supplementary File 2). Oligos used to construct plasmids are listed in Table S1. Plasmids used to knock-in *promoter::TIR1::F2A::BFP::AID*::NLS* sequences are listed in Table S2. All other plasmids are listed in Table S3. Primer design information for designing homology arms for Gibson and SapTrap cloning is provided in Supplementary File 3.

### C. elegans

*C. elegans* was cultured as originally described (Brenner 1974). The majority of genome editing was performed in N2 (wild type), EG9615 {oxSi1091[*mex-5p::Cas9(smu-2 introns) unc-119+] II; unc-119(ed3) III}*, or EG9882 (*F53A2*.*9{oxTi1127[mex-5p::cas9(+smu-2 introns)]}, hsp-16*.*41p::Cre, Pmyo-2::2xNLS-CyOFP + lox2242 III)* animals (Table S4). EG9615 and EG9882 (unpublished) stably express Cas9 in the germline and are gifts from Dr. Matthew Schwartz and Dr. Erik Jorgensen. The *mex-5p* and *cdh-3p* strains were generated in specialized genetic backgrounds and then the TIR1 transgene was removed by outcrossing. Details are provided in Supplemental File 2. Strains expressing TIR1 generated through genome editing are listed in the table in Figure 3A. We note that we are reporting the final, SEC-excised strains in this table. We will also make the precursor strains containing the SEC available to the CGC for our *mex-5*p and *eft-3p* strains: JDW220 *wrdSi10[mex-5p::TIR1::F2A::BFP::tbb-2 3’UTR+SEC] I*, JDW222 *wrdSi8[mex-5p::TIR1::F2A::BFP::tbb-2 3’UTR + SEC] II*, and JDW224 *wrdSi22[eft-3p::TIR1::F2A:BFP::tbb-2 3’UTR+SEC] I*.

**Figure 3.**
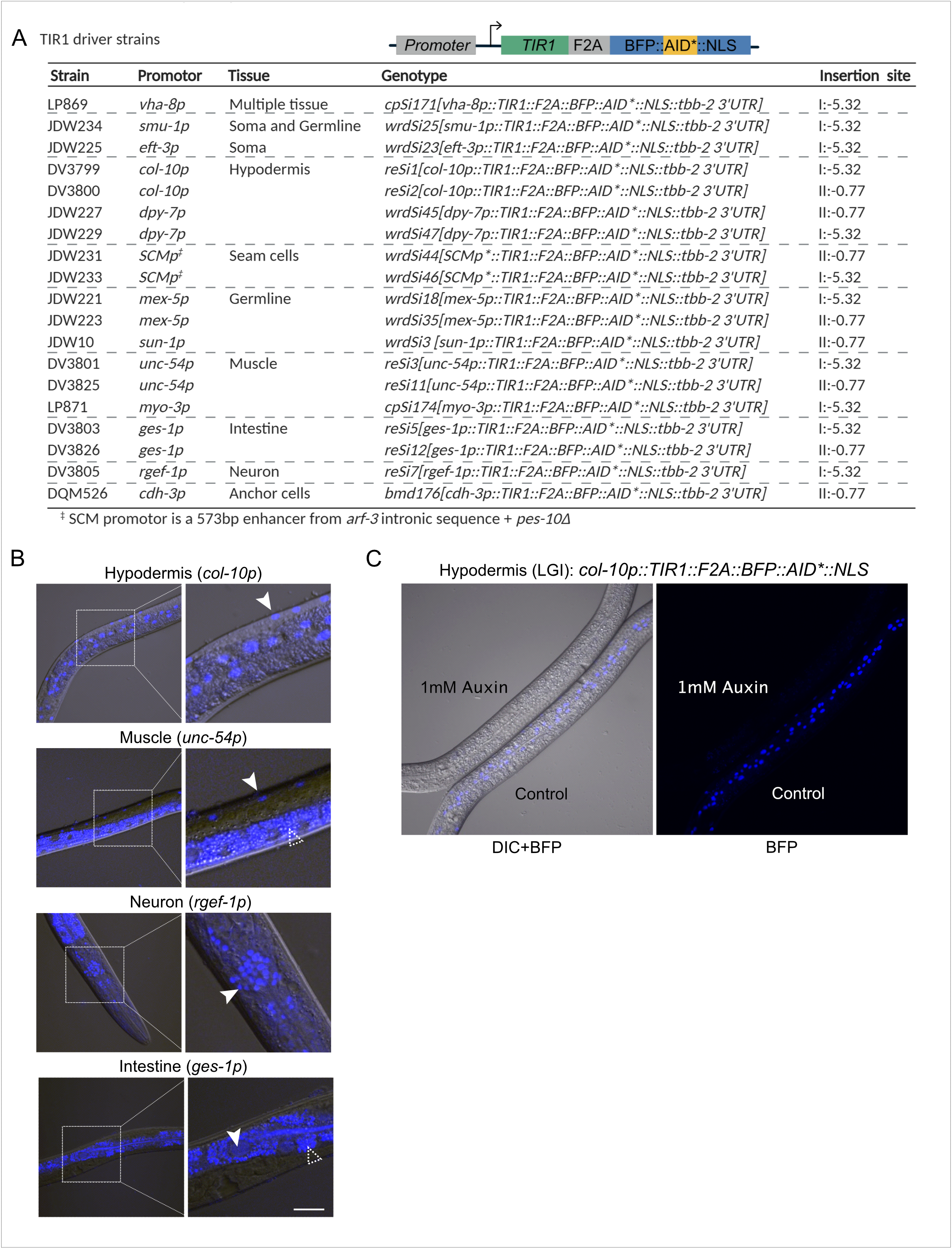
A new suite of TIR1 expression strains for tissue-specific depletion of AID-tagged proteins in *C. elegans*. A) Table describing new suite of TIR1::F2A::BFP::AID*::NLS strains. Strain names, promoter driving TIR1, tissue of expression, genotype, and insertion site are provided for each strain. The insertion sites are the genomic loci where the MosI transposon landed in the ttTi4348 and ttTi5605 insertion alleles. We note that our knock-ins were generated using CRISPR/Cas9-mediated genome editing in wildtype animals or in strains stably expressing Cas9 in the germline; there is no MosI transposon in these loci in these genetic backgrounds. B) BFP is detected in the expected nuclei of strains expressing TIR1 cassettes driven by *col-10p* (hypodermis), *unc-54p* (muscle), *ges-1p* (intestine), and *rgef-1p* (neurons). A representative BFP expressing nucleus is indicated by a solid arrow. Scale bars represent 20 µm. Note that the fluorescence signal at the bottom of the muscle image and surrounding the nuclei in the intestinal image is intestinal autofluorescence, and indicated by an unfilled arrow with a dashed outline. C) Functional test of TIR1 activity in a *col-10p::TIR1::F2A::BFP::AID*::NLS* strain (DV3799). Hypodermal BFP expression is lost when animals are exposed to 1 mM auxin for three hours, but not when similarly grown on control plates.

### CRISPR/Cas9-based genome editing

All TIR1 strains were generated through SEC selection-based genome editing as previously described (Dickinson *et al*. 2015). Single-copy transgenes were inserted into the chromosome I and II loci where the ttTi4348 and ttTi5605 transposons are inserted for MosSCI (Frøkjaer-Jensen *et al*. 2008; Frøkjær-Jensen *et al*. 2012), respectively. The genetic map positions for these insertions are provided in the genotype information. In Table S4, we detail the genetic background in which injections were performed, which Cas9 and sgRNA plasmids were used, and how many times the strains were outcrossed against an N2 background. Repair templates were used at 10 ng/µl and Cas9+sgRNA or sgRNA plasmids were used at 50 ng/µl.

### Auxin treatment

For DQM623 (Figure 4) and JDW221, JDW225, and JDW229 (Figure S1), one-hour auxin treatments were performed. Worms were synchronized at the L1 larval stage by sodium hypochlorite treatment and moved to nematode growth media (NGM) plates seeded with *E. coli* OP50. At the young adult stage, JDW221 worms were either kept on OP50-seeded NGM plates (control) or moved to plates treated with 1 mM K-NAA (Phyto-Technology Laboratories, N610) (Martinez and Matus 2020). At the mid-L3 larval stage, JDW225 and JDW229 worms were either kept on OP50-seeded NGM plates or moved to plates treated with 1 mM K-NAA. At the early L3, DQM623 worms were either kept on OP50-seeded NGM plates or moved to plates treated with 4 mM K-NAA. OP50-seeded NGM plates containing K-NAA were prepared as described (Martinez and Matus 2020). For LP869 and LP871, mixed stage animals were transferred to 1 mM K-NAA NGM plates for 24 hours and imaged as described.

**Figure 4.**
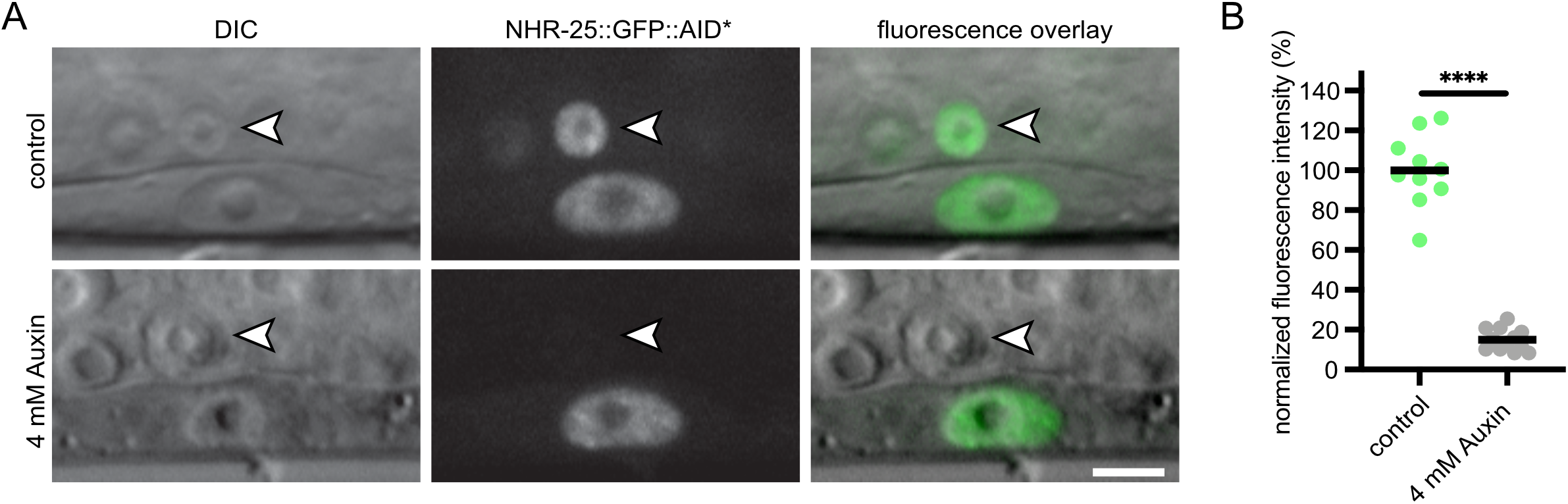
NHR-25::GFP::AID*::3xFLAG can be depleted in a cell-specific manner in a strain with undetectable TIR1 expression via a BFP reporter. A) An anchor cell (AC)-specific TIR1 transgene (*cdh-3p::TIR1::F2A::BFP::AID*::NLS*) did not produce observable BFP in the AC. Crossing this strain to an *nhr-25::GFP::AID*::3xFLAG* allele resulted in depletion of NHR-25 in the AC when exposed to 1 mM Auxin for 1 hr (indicated by white arrow with black outline). Depletion of NHR-25 was not observed in the neighboring uterine cells or the underlying vulval precursor cells (VPCs). Scale bar represents 5 µm. B) Quantification of NHR-25::GFP::AID*::3xFLAG in ACs following auxin (K-NAA) treatment. Data presented as the mean (n ≥ 10 animals examined for each condition; **** indicates *P* < 0.0001 by a two-tailed unpaired Student’s t-test. *P* < 0.05 was considered statistically significant). Scale bars represent 5 µm.

For DV3799, DV3801, DV3803, and DV3805 (Figure 3), we made an auxin (IAA) (Alpha Aesar, #A10556) stock solution (400 mM in ethanol) and stored at 4ºC for up to one month. A 16 mM auxin working solution was then prepared freshly by diluting 1:25 in filtered water with 4% ethanol final concentration. Animals were cultured on OP50 seeded on 60 mm NGM plates to the required stage, and 500 µL of 16 mM auxin was added to plate for a final concentration of 1 mM, with 4% ethanol as vehicle control (plates contain approximately 8 ml of agar). Animals were treated with auxin or vehicle for 3 hours before imaging.

### Microscopy

For DQM623 in Figure 4 and JDW221, JDW225, and JDW229 in Figure S1, images were acquired on a custom-built upright spinning-disk confocal microscope consisting of a Zeiss Axio Imager.A2, a Borealis-modified Yokogawa CSU10 confocal scanner unit with 50 mW, 405 nm lasers and 25 mW, 488 nm lasers, and a Hamamatsu Orca EM-CCD camera. Images shown for JDW221 (pachytene region) were acquired using a Plan-Apochromat 40x/1.4 DIC objective. Images shown for DQM623 (AC), JDW225 (uterine and vulval tissues) and JDW229 (hypodermal cells) were acquired using a Plan-Apochromat 100x/1.4 DIC objective. MetaMorph software (version: 7.8.12.0) was used to automate acquisition. Worms were anesthetized on 5% agarose pads containing 7 mM NaN_3_ and secured with a coverslip. Acquired images were processed through Fiji software (version: 2.0.0-rc-69/1.52p).

The LP869 and LP871 images (Figure S1) were taken using the 60x objective on a Nikon TiE stand with CSU-X1 spinning disk head (Yokogawa), 447 nm, 514 nm, and 561 nm solid state lasers, ImagEM EMCCD camera (Hamamatsu). Worms were anesthetized and images were processed as described above.

For strains DV3799, DV3800, DV3801, DV3803, DV3805, DV3825, and DV3826 (Figure 3), animals were anaesthetized with 5 mM tetramisole. Images were acquired on a Nikon A1si Confocal Laser Microscope using a Plan-Apochromat 40x/1.4 DIC objective and DS-Fi2 camera. Images were analyzed using NIS Elements Advanced Research, Version 4.40 software (Nikon).

### Statistical analysis

Statistical significance was determined using a two-tailed unpaired Student’s t-test. *P* < 0.05 was considered statistically significant. \*\*\*\**P* < 0.0001. The graph in Figure 4B was made using Prism software (version: 8.4.2.).

#### Data availability

Strains in Figure 3A will be made available through the *Caenorhabditis* Genetics Center. Other strains and plasmids can be requested directly from the authors. The data that support the findings of this study are available upon reasonable request. pDD356 and pDD357 as well as plasmids described in Figures 5 and 6 will be made available through Addgene. Supplemental files will be available on figshare.

**Figure 5.**
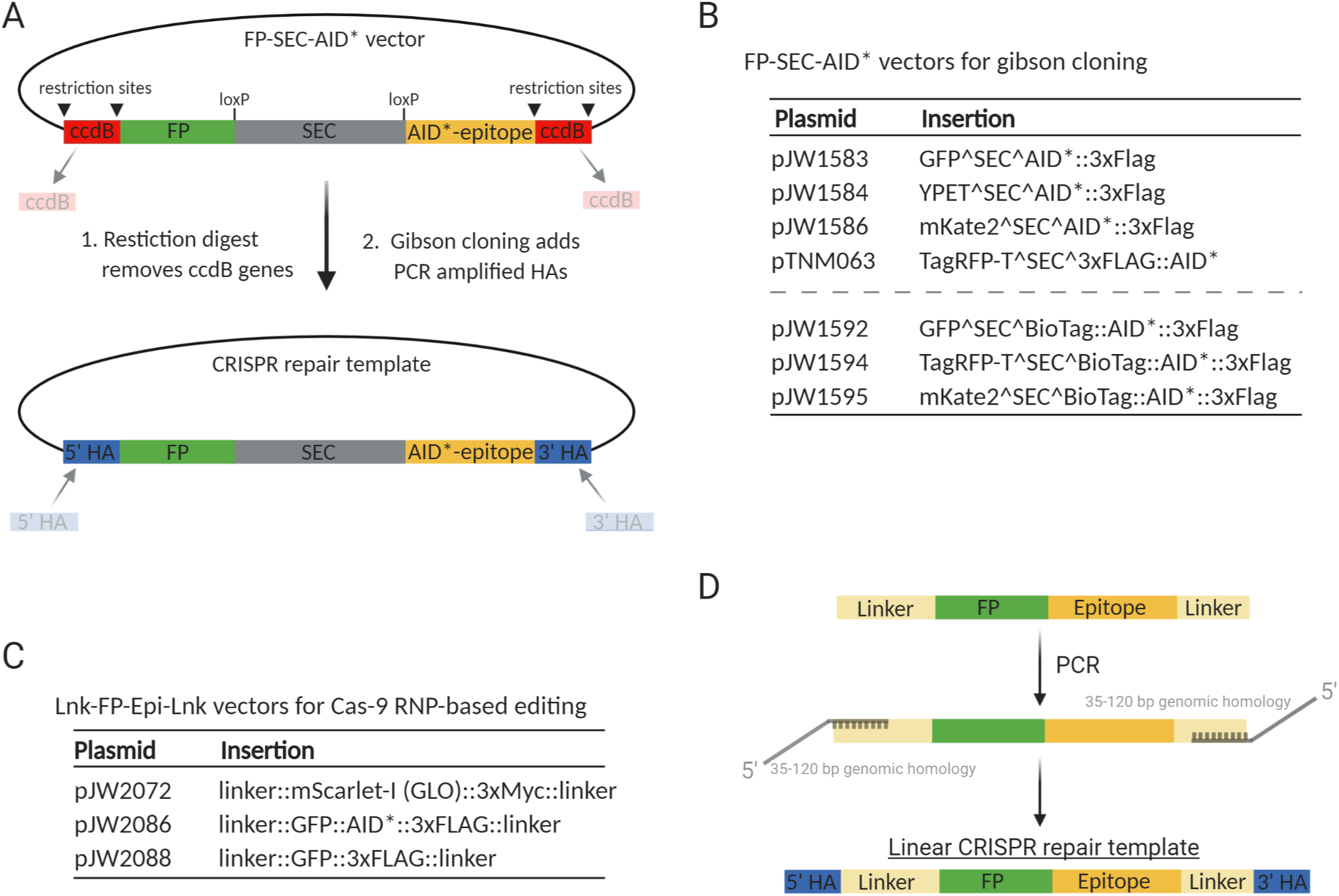
A collection of vectors to generate FP::AID* knock-ins through Gibson cloning into self-excising cassette (SEC) vectors or through generation of linear repair templates. A) Schematic of the AID* containing vectors produced by modifying the set of vectors originally described by Dickinson *et al*. (2015). An AID* epitope was inserted downstream of the loxP-flanked SEC. New repair templates for CRISPR/Cas9-mediated genome editing are produced by restriction digestion of the vector and Gibson cloning of PCR-derived homology arms, as described (Dickinson *et al*., 2015). Counterselection against the parent vector is provided by ccdB cassettes. B) Suite of FP::AID* SEC vectors available through Addgene. The vectors described in Dickinson *et al*. (2015) have been modified to insert an AID* or 23 amino acid biotin acceptor peptide (BioTag)::AID* cassette between the SEC and 3xFLAG cassette. C) A set of vectors to generate repair templates for Cas9 ribonucleoprotein complex (RNP)-based genome editing. FP and FP::AID* cassettes are flanked by flexible linker sequences. A 30 amino acid sequence is at the 5’ end of the cassette, and a 10 amino acid sequence is at the 3’ end of the cassette. This design provides flexibility for designing repair templates for N-terminal, C-terminal, or internal tagging. D) Schematic of how to generate linear repair templates by PCR. Primers with homology to the cassette and 5’ homology to the desired integration site are used to amplify a dsDNA repair template. 35-120 bp homology arms are recommended, as previously described (Paix *et al*. 2014; 2015; Dokshin *et al*. 2018).

**Figure 6.**
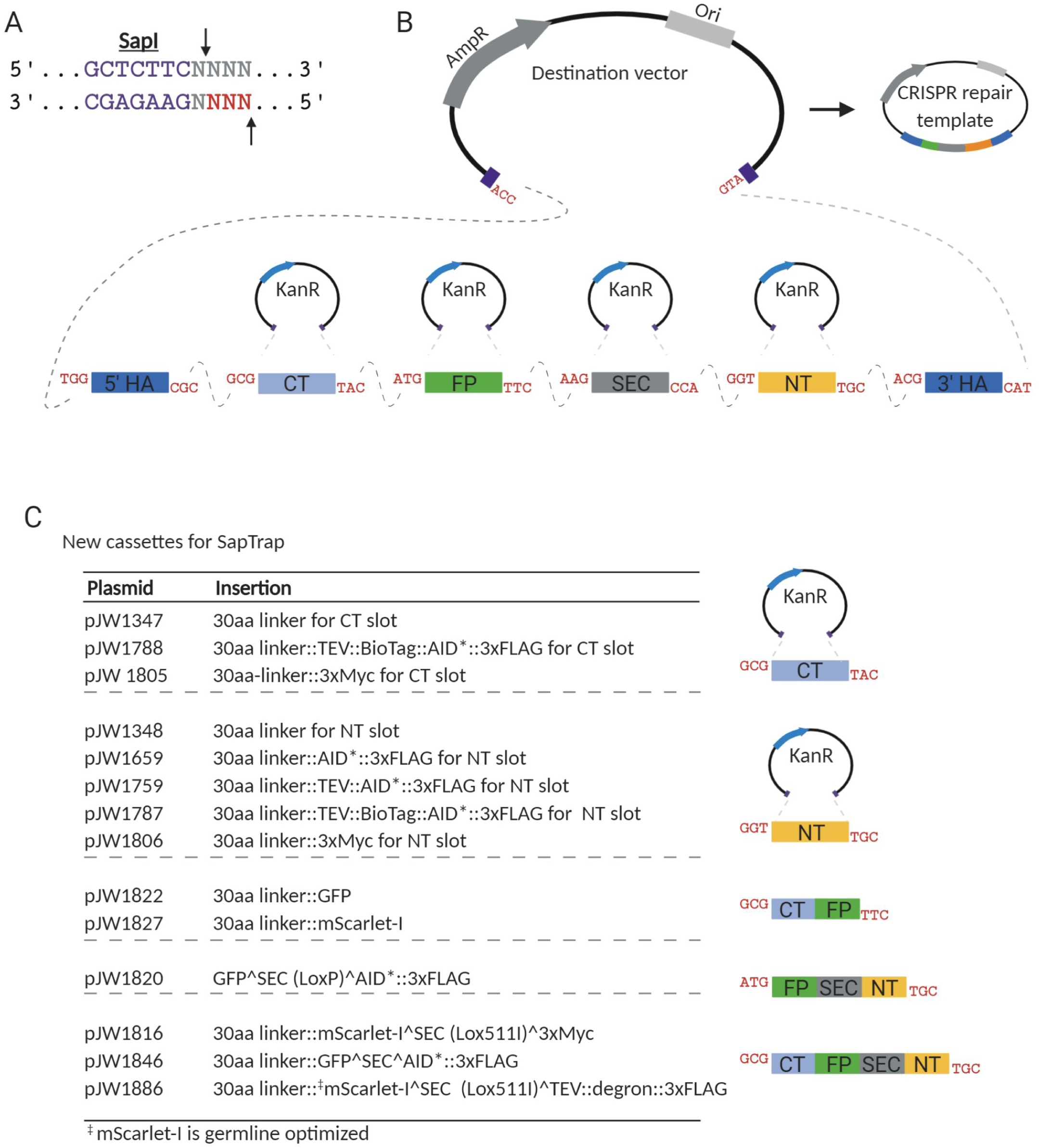
A suite of new vectors for the SapTrap cloning system. A) SapI is a type II restriction enzyme that cuts one base pair and four base pairs outside of its binding site, allowing for the generation of programable 3bp sticky ends. B) SapTrap cloning facilitates single reaction cloning of multiple fragments, in the correct order, into a single repair template plasmid. Specific sticky ends are used for specific cassettes. C) Table of new vectors generated for the SapTrap CT and NT slots. Our initial assembly efficiencies were sub-optimal, and we found that reducing the number of fragments assembled improved our efficiencies. We have generated a set of multi-cassettes where partial assemblies (CT-FP, FP-SEC-NT, and CT-FP-SEC-NT) have been cloned, simplifying the SapTrap reactions and reducing the number of fragments required.

## Results

### A new suite of TIR1 driver strains compatible with red FP imaging

The initial description of the AID system in *C. elegans* used an mRuby2 fusion to monitor the expression of TIR1 (Zhang *et al*. 2015). This feature was useful to monitor TIR1 localization and expression level in comparison to depletion of GFP::AID* tagged proteins. One limitation is that the TIR1::mRuby2 interferes with the imaging of factors tagged with red FPs. Blue fluorescent proteins (BFPs) offer an appealing alternative as reporters for TIR1 expression, since their emission spectra do not overlap with commonly used green and red FPs (Lambert 2019). To report TIR1 localization and activity, we placed an *F2A::BFP::AID*::NLS* reporter downstream of the TIR1 transgene and drove germline and embryo expression with a *sun-1* promoter (Figure 2). F2A is an example of a 2A peptide, a virally derived ribosome skip sequence that allows production of multiple polypeptides from a single mRNA (Ryan and Drew 1994; Ryan *et al*. 1999; Donnelly *et al*. 2001; de Felipe *et al*. 2003). These sequences function effectively in *C. elegans* and allow up to five proteins to be produced from a single mRNA (Ahier and Jarriault 2014). Transgenes were introduced in single copy through CRISPR/Cas9 editing and self-excising cassette (SEC) selection into neutral loci that support robust expression. We chose the sites in chromosomes I and II, respectively, where the ttTi4348 and ttTi5605 transposons are inserted for MosSCI-based genome editing (Frøkjaer-Jensen *et al*. 2008; Frøkjær-Jensen *et al*. 2012). These vectors are a useful counterpart to the MosSCI vectors; they allow introduction of transgenes into strains lacking Mos1 transposons, and thus can be inserted in any strain. The SEC strategy (Dickinson *et al*. 2015) first produces hygromycin resistant, rolling animals, which is useful for tracking the allele phenotypically in crosses. The loxP-flanked SEC is then excised by heat shock, producing the final wildtype-moving strain. As expected, this *sun-1p* construct drives nuclear-localized BFP in the germline and embryos, confirming the expression of the transgene (Figure 2B). We confirmed TIR1 activity by placing adult animals on 1 mM auxin and observing loss of BFP::AID*::NLS (Figure 2B). We attempted to generate an equivalent construct where the *TIR1::F2A::BFP::AID*::NLS* sequence was codon optimized and had piRNA sites removed, as this has been shown to improve expression of germline transgenes (Zhang *et al*. 2018a). Although we successfully generated single-copy insertions of this construct, we did not observe detectable expression of the piRNA-depleted TIR1::F2A::BFP::AID*::NLS protein in any animals from three independent transgene insertion lines. We have not pursued whether these issues are due to transgene toxicity, silencing, or other potential issues.

Using our new TIR1 construct, we created a suite of strains with ubiquitous or tissue-specific TIR1 expression (Figure 3A). We created chromosome I and II knock-ins expressing TIR1 in the germline (*mex-5p* and *sun-1p*), hypodermis (*dpy-7p* and *col-10*p), muscle *(unc-54p*), and intestine (*ges-1p)* (Figure 3A, Figure S1). We also created chromosome I knock-ins expressing TIR1 in neurons (rgef-1p), somatic cells (eft-3p), body wall muscle (myo-3p), and excretory cell+hypodermis+gut (vha-8p) (Figure 3A, Figure S1). Our vha-8p strain also resulted in promoter expression in unidentified cells in the head. We also generated a strain expressing TIR1 in the seam cells using a minimal SCMp enhancer (gift from Prof. Allison Woollard) and a pes-10 minimal promoter (Figure 3A). While we saw robust seam cell expression in this strain, we also detected hypodermal expression (unpublished data). We are making this strain available to the community, but encourage careful evaluation before interpretation.

An unanswered question is the importance of TIR1 expression levels for effective depletion of AID*-tagged proteins. Motivated by an interest in NHR-25 (Ward *et al*. 2013; 2014) and anchor cell invasion (Matus *et al*. 2015; Medwig and Matus 2017; Medwig-Kinney *et al*. 2020), we generated a strain to facilitate anchor cell (AC)-specific protein depletion (DQM623; Figure 3A). We observed no detectable BFP in this *cdh-3p::TIR1::F2A::BFP::AID*::NLS* strain. To perform a functional test, we crossed in an *nhr-25::GFP::AID*::3xFLAG* allele into the *cdh-3p::TIR1*. We had previously demonstrated significant depletion of NHR-25 in ACs and vulval precursor cells (VPCs) using a strongly expressed *eft-3p::TIR1::mRuby2* transgene (Martinez *et al*. 2020). Strikingly, we observed auxin-dependent depletion of NHR-25::GFP::AID*::3xFLAG in the AC, while no depletion was observed in the adjacent VPCs (Figure 4). Thus, even if the presence of TIR1 is undetectable through BFP reporter expression, there may still be sufficient amounts of TIR1 to deplete proteins of interest. We also made a strain designed to express TIR1 in both the soma and germline (smu-1p), as the eft-3p driven TIR1 transgenes are typically silenced in the germline. We could not detect BFP expression in this smu-1p strain, but have made it available for the community to test. The majority of the TIR1 strains (17/19) we will deposit in the Caenorhabditis Genetics Center have detectable BFP expression that is lost when animals are shifted onto auxin plates, confirming TIR1 is active (Figures 2, 3 and Figure S1); the exceptions to this statement are the previously discussed *smu-1p* and *cdh-3p* TIR1 strains.

### Vectors to generate FP::AID* knock-ins

Currently, there are few vectors in repositories such as Addgene that allow facile customization of FP::AID* repair templates for generating knock-ins into endogenous genes using CRISPR/Cas9-mediated genome editing. Zhang *et al*. (2015) described *unc-119* selectable linker::AID*::GFP vectors, and Dickinson *et al*. (2018) generated two AID* cassettes for the SapTrap system. We, therefore, created a set of vectors compatible with the most common genome-editing pipelines we use in our lab. First, we took a set of vectors that use Gibson assembly to generate SEC-selectable repair templates developed by Dickinson *et al*. (2015) and introduced AID* sequences upstream of the 3xFLAG epitope. This set of vectors allows for tagging genes with GFP, YPET, mKate2, and TagRFP-T along with AID*::3xFLAG epitopes (Figure 5, Table S3). Methods in *C. elegans* using biotin ligases for protein affinity purification (Waaijers *et al*. 2016), proximity labeling (Branon *et al*. 2018), native chromatin purification (Ooi *et al*. 2010), and cell-type specific nuclei purification (Steiner *et al*. 2012) have recently been developed. To support these approaches, we have made a set of FP^SEC^BioTag::AID*::3xFLAG vectors. The BioTag sequence was the same as was used by Ooi *et al*. (2010) and Steiner *et al*. (2012), and we constructed vectors with GFP, TagRFP-T, and mKate2.

We also have shifted to frequently using Cas9 RNP-based editing with linear repair templates, as this approach is cloning-free (Paix *et al*. 2014; 2015), and recent refinements have further boosted editing efficiency (Dokshin *et al*. 2018). We have generated plasmids with linker::GFP::AID*::3xFLAG::linker (pJW2086), linker::GFP::3XFLAG::linker (pJW2088), and linker::mScarlet-I (germline-optimized)::3xMyc::linker (pJW2072) cassettes (Figure 5, Table S3). These vectors are suitable templates for PCR amplification to generate linear repair templates with short homology arms. FPs can be easily exchanged to generate new constructs by PCR linearization and Gibson cloning. Our standard approach is to include the AID* tag on a GFP-tagged protein and examine the impact of depletion of this protein on a second mScarlet-I::3xMyc-tagged protein. Linkers flanking the FP allow flexibility in targeting genes of interest and reduce functional interference, permitting N-terminal, C-terminal, or internal tagging.

Modifying the large, SEC-based selection cassettes for Gibson assembly was technically challenging as the size and repetitive *ccdB* sequences made these prone to recombination. The modularity of the type II restriction enzyme-based SapTrap cloning pipeline was appealing as a method to rapidly develop new repair templates for CRISPR/Cas9-mediated genome editing (Figure 6A). Recent modifications to the system have made it compatible with SEC-based cloning (Dickinson *et al*. 2018), retaining an FP cassette and adding an SEC selection cassette. We generated a series of constructs for the SapTrap NT and CT slots, consisting of flexible linkers, various combinations of AID* cassettes, and epitopes for protein purification or detection (3xMyc, 3xFLAG, BioTag (Figure 6B). We first attempted to generate a 30 amino acid linker::GFP^SEC^TEV::AID*::3xFLAG construct to tag *lin-42* at the C-terminus. While we were able to produce the construct and obtain a knock-in strain (unpublished), our efficiency was very low, and we were unable to get correct assemblies by simply selecting colonies and sequencing. We, therefore, turned to colony PCR screening. Our standard practice of screening one assembly junction produced false positives, where one homology arm was correctly connected to the backbone and desired SapTrap cassettes, but the other arm was missing cassettes. We therefore screened both assembly junctions with the vector by colony PCR to identify correct clones, finding one correct assembly out of 48 colonies. As this efficiency was much lower than reported in the original SapTrap description (Schwartz and Jorgensen 2016), we obtained the reagents for generating an *snt-1::GFP* targeting vector (gift from Dr. Matt Schwartz and Dr. Erik Jorgensen). We obtained a similar assembly efficiency as reported (Schwartz and Jorgensen 2016), indicating that our efficiency issues were not due to our SapTrap reagents. In examining our colony PCR data for the *lin-42* construct, we occasionally noted the presence of bands smaller than the expected product. Sequencing the plasmid from these strains revealed partial assemblies of 2-3 blocks. We then PCR-amplified these partial assemblies to restore the terminal SapI sites and connectors. Using these partial assembly blocks dramatically improved efficiency. To facilitate an SEC-based SapTrap assembly pipeline, we generated a series of “multi-cassettes,” where we combined fragments that we frequently use (Figure 6C, Table S3). We re-created the 30 amino acid linker::GFP^SEC^TEV::AID*::3xFLAG *lin-42* targeting construct using a multi-cassette, and our colony PCR hit rate jumped from 1/48 (2.1% (70.8%)) to 17/24. For our most commonly used vectors, we have generated constructs containing full assemblies of the knock-in epitope, lacking only the homology arms. PCR amplifying homology arms with SapI sites and appropriate connectors allows high-efficiency generation of repair templates.

### Additional vectors to support genome editing and gene expression studies

We had previously shown that the “Flipped and extended (F+E)” sgRNA modifications that improved editing efficiency in mammalian cells (Chen *et al*. 2013) had a similar impact in *C. elegans* (Ward 2015). This modification was recently introduced into a SapTrap repair template vector that also contains a *U6p::sgRNA* cassette (Dickinson *et al*. 2018). We frequently used a separate sgRNA vector to reduce the number of fragments in an assembly and to allow us to increase the molar ratio of sgRNA vector to repair template. We generated SapTrap sgRNA RNA expression vectors using both commonly used U6 promoters (pJW1838,pJW1839)), as it is currently unclear whether one promoter is broadly more active, (Friedland *et al*. 2013; Dickinson *et al*. 2013; Schwartz and Jorgensen 2016). One study reported that the K09B11.2 U6 promoter produces higher editing frequencies for a *sqt-1(sc1)* knock-in (Katic *et al*. 2015), while another study was unable to generate *rol-6(su1006)* alleles using the K09B11.2 U6 promoter, but, instead, had success with the R07E5.16 promoter (Farboud and Meyer 2015)(Table S3). Our TIR1 transgenes were inserted into the same standardized chromosomal loci frequently used for MosSCI-based generation of transgenes (ttTi4348 and ttTi5605) (Frøkjaer-Jensen *et al*. 2008; Frøkjær-Jensen *et al*. 2012). We therefore created sgRNA(F+E vectors) for both U6 promoters for these two loci, as well as a third standardized locus (cxTi10882 insertion site) (pJW1849-1851; pJW1882-1884)(Frøkjær-Jensen *et al*. 2012). We have also generated vectors containing *eft-3p::Cas9*+*R07E5*.*16 U6p::sgRNA (F+E)* (Dickinson *et al*. 2013; Ward 2015) targeting the ttTi4348 and ttTi5605 insertion sites (pTD77 and pTD78, respectively). Finally, a set of germline optimized (Redemann *et al*. 2011; Wu *et al*. 2018) NLS::mScarlet-I vectors used as a cloning intermediate provide useful promoter reporter constructs. These Gibson-cloneable vectors come in versions that are promoterless (pJW1836), or have a *pes-10Δ* minimal promoter (pJW1841) for testing enhancers. We also made destabilized versions of these reporters by adding a PEST sequence (Loetscher *et al*. 1991), to allow monitoring of dynamic promoter activity (pJW1947,1948).

## Discussion

The AID system has allowed rapid, conditional, and tissue-specific depletion of tagged proteins in a wide range of organisms and cell types. Since its introduction to *C. elegans* (Zhang *et al*. 2015), it has been promptly adopted by the community. This system has allowed for rapid depletion of proteins in tissues that are refractory to RNA interference approaches, such as the germline (Pelisch *et al*. 2017; Shen *et al*. 2018; Zhang *et al*. 2018b), vulval precursor cells (Matus *et al*. 2014), and neurons (Liu *et al*. 2017; Patel and Hobert 2017; Serrano-Saiz *et al*. 2018). The system is also powerful for studying rapid developmental events such as molting (Zhang *et al*. 2015; Joseph *et al*. 2020), organogenesis (Martinez *et al*. 2020), and developmental timing (Azzi *et al*. 2020). Improvements to the auxin ligand have enhanced protein degradation in the embryo (Negishi *et al*. 2019) and removed the need for ethanol solubilization, instead allowing the auxin derivative to be dissolved in any aqueous buffer (Martinez *et al*. 2020). This soluble auxin was shown to be compatible with microfluidic devices, allowing long-term imaging coupled with targeted protein depletion (Martinez *et al*. 2020). Auxin-mediated depletion of a spermatogenesis regulator has been developed to conditionally sterilize animals, a valuable approach for the *C. elegans* aging field (Kasimatis *et al*. 2018). Given the emerging use of the AID system in *C. elegans*, it is important to continue to develop strains and reagents to facilitate usage.

The original *C. elegans* AID system employed a TIR1::mRuby2 transgene, which was useful for visualizing TIR1 expression and cellular localization (Zhang *et al*. 2015). However, for applications where red fluorescent protein imaging is desired, the mRuby2 expression could increase background and hamper imaging analysis. We therefore developed a complementary construct containing a *TIR1::F2A::BFP::AID*::NLS* transgene (Figure 2). TIR1 is unlabeled, and nuclear-localized BFP provides an equimolar readout for TIR1 expression (Figure 2). AID*-tagged BFP is degraded in the presence of auxin, confirming TIR1 activity via degradation of an internal control (Figure 2). The ability to read out TIR1 activity opens the door to performing suppressor screens for phenotypes of interest generated using the AID system. Mutations in the TIR1 transgene or auxin transport factors could lead to unintended suppression of a mutant phenotype when performing such a screen. Therefore, being able to monitor TIR1 activity provides a secondary screen for such mutations. The vectors containing the *TIR1::F2A::BFP::AID*::NLS* transgene are SEC-based vectors targeting the ttTi4348 and ttTi5605 insertion sites. These vectors contain unique restriction sites to remove promoters or even the entire transgene, facilitating creation of new TIR1 drivers or transgenes by Gibson assembly. The dominant *sqt-1(sc13)* marker in the SEC is useful for tracking loci during crosses via a roller phenotype and the SEC can easily be excised by heat shock, as previously described (Dickinson *et al*. 2015).

We have also generated vectors compatible with commonly used cloning and genome editing protocols to facilitate knock-in of FP::AID* constructs into genes of interest (Figures 5,6, Table S3). The set of Gibson-cloneable SEC vectors originally described by Dickinson *et al*. (2015) have been widely used, so we created modified versions containing AID* and BioTag::AID* cassettes (Figure 5). While we have not extensively tested the BioTag, it will be useful for streptavidin-based purification protocols developed by the community (Steiner *et al*. 2012; Steiner and Henikoff 2014; Waaijers *et al*. 2016; Branon *et al*. 2018). The strength of these vectors is that they provide easy, highly efficient assembly of repair templates for CRISPR/Cas9-mediated genome editing. One challenge in generating new versions of these vectors containing new fluorescent proteins or epitopes is that they are large and have repetitive sequences, making them prone to rearrangement. We turned to SapTrap, as the modular design and ligation-based cloning should simplify integration of new FPs and epitopes. As we typically perform edits in a range of genetic backgrounds, we designed our constructs to use the SEC cassettes designed for the SapTrap system (Dickinson *et al*. 2018). In our first tests requiring assembly of nine pieces of DNA (two homology arms, sgRNA, four cassettes, two plasmid backbone fragments), we had poor efficiency and required screening of two assembly junctions by colony PCR to identify a single correct assembly out of 48 colonies. While the SEC-based version of SapTrap was less efficient than Gibson cloning (∼60% vs. 30%) (Dickinson *et al*. 2018), it was still much higher than the efficiencies we initially observed. In screening our reactions, we would occasionally identify partial assemblies (i.e. 5’ homology arm+CT cassette). By cloning out these partial assemblies to reduce the number of fragments, we boosted our assembly efficiencies. We then shifted to create multi-cassettes, which minimize the number of ligations required to successfully assemble the final vector. Additionally, by expressing our sgRNA on a separate plasmid, only four pieces of DNA need to be assembled in all, boosting efficiency (backbone, 5’ homology arm, 3’ homology arm, multi-cassette). We also provide methods for the community to build new multi-cassettes for highly used FP-epitope combinations. Finally, we generated a set of vectors for making linear repair templates by Cas9 RNPs (Figure 5). The advantage of this approach is that it does not require cloning and has high reported efficiencies (Paix *et al*. 2014; 2015; Dokshin *et al*. 2018). These vectors contain N- and C-terminal flexible linkers so that they can be used for tagging proteins of interest at the N-terminus, C-terminus, or internally. The small size of these plasmids allows FPs and epitopes to easily be exchanged by Gibson cloning. Together, this collection of vectors should facilitate efficient generation of new *FP::AID\**-tagged genes.

For many applications, the AID system offers a powerful method to conditionally degrade proteins in specific tissues and at specific points in development. However, as the system has gained popularity, particular challenges have emerged. While they do not dampen our enthusiasm for the AID system, it is important to be aware of them. Here, we also discuss potential solutions to these issues. It has become clear that in certain cases there can be auxin-independent, TIR1-dependent degradation of AID-tagged proteins. This unwanted basal degradation occurs with both the minimized and full-length AID sequence. In mammalian cells, a recent report found 3-15% depletion of AID-tagged proteins in the absence of auxin when TIR1 is co-expressed (Sathyan *et al*. 2019). This auxin-independent degradation can be extreme: in human cell lines, tagging the centromeric histone chaperone HJURP with AID resulted in over 90% depletion in the absence of auxin (Zasadzińska *et al*. 2018). In line with these findings, in *C. elegans* RNAi depletion of the Cullin-1 gene, *cul-1*, increased expression of a GFP::AID reporter by 19% in the absence of auxin and presence of TIR1 (Martinez *et al*. 2020). Martinez *et al*. (2020) also reported 22-35% depletion of AID-tagged *C. elegans* proteins in an auxin-independent, TIR1-dependent manner, while another study observed 70-75% depletion (Schiksnis *et al*. 2020). Currently, the determinants of a protein’s sensitivity to auxin-independent degradation are unclear. Sathyan *et al*. (2019) reported that the addition of another component of the auxin signaling pathway, ARF19, blocked this auxin-independent degradation of AID-tagged proteins and in fact sped up degradation kinetics in the presence of auxin. ARF19 is an AID interaction partner and is thought to shield the tagged protein from TIR1 in the absence of auxin. It may be useful to test whether ARF19 improves performance of the AID system in *C. elegans*. One important caveat is that the authors used a full-length AID tag. The miniAID and AID* tags frequently used in *C. elegans* lack domains III and IV of the protein which are thought to be important for the ARF19 interaction. Full-length AID is 229 amino acids, a substantially larger tag that would necessitate further study to ensure it did not interfere with fusion proteins. Another approach could be a recently described AID system comprised of *Arabidopsis thaliana* AFB2 and a minimal degron from IAA7, which was reported to minimize basal degradation (Li *et al*. 2019). Our set of vectors will allow modular assembly of any new AID system component and facile integration of any new reagents. We note that engineering an improved TIR1 that did not promote auxin-independent degradation of miniAID-tagged proteins would be most desirable. A strong candidate is a recently described TIR1 (F79A) mutation and modified auxin that had 1000-fold stronger binding, reducing the amount of auxin required for target knockdown (Nishimura *et al*. 2020). This reagent would be compatible with and improve the performance of the collection of miniAID- and AID*-tagged strains which the *C. elegans* community has already generated.

We previously used the AID system to deplete the nuclear hormone receptor NHR-23 and reported a larval arrest phenotype similar to a previously-described null allele, and depletion of NHR-23 within 20 minutes of auxin exposure (Kouns *et al*. 2011; Zhang *et al*. 2015). This result highlights the potential of the AID system for rapid depletion of proteins of interest. However, AID-mediated protein depletion may not always produce strong hypomorphic or null phenotypes. Serrano-Saiz *et al*. reported that with an *unc-86::mNeonGreen::AID* allele combined with ubiquitously-expressed TIR1 via a strong *eft-3* promoter, and continuous exposure to 4 mM auxin, they failed to observe the phenotype of strong loss-of-function or null alleles. Despite a complete loss of mNeonGreen expression via confocal microscopy, they only observed phenotypes consistent with an *unc-86* hypomorph (Serrano-Saiz *et al*. 2018). In interpreting the extent of depletion by microscopy, one must consider the limit of detection and fluorophore maturation time. In adult males, a GFP::AID*::3xFLAG tagged allele of NHR-23 is only detectable in the germline (manuscript in preparation). When we use a germline-specific TIR1 to deplete NHR-23, by microscopy we observed no detectable expression following auxin exposure (manuscript in preparation). However, ∼30% of protein remains as detected by western blot (manuscript in preparation). A similar inability to obtain null phenotypes using the AID system was observed using an *unc-3::mNeonGreen::AID\** (Patel and Hobert 2017) allele and a *daf-15::mNeonGreen::AID\** allele (Duong *et al*. 2020). In addition, the mNeonGreen::AID* tag caused a mild hypomorph of *unc-3* in the absence of TIR1 and auxin, suggesting that the presence of the AID* tag was interfering with protein levels (Patel and Hobert 2017). The inability of target protein depletion to produce a null phenotype has been encountered for several other neuronal identity genes (Oliver Hobert, personal communication). In yeast, fusion of the Skp1 subunit of the SCF ubiquitin ligase to TIR1 has resulted in enhanced degradation efficiency of AID-tagged proteins (Kanke *et al*. 2011). Such an approach is worth testing in *C. elegans*. More examples are needed to assess the likely mechanism of TIR1-independent, auxin-independent inactivation of target proteins. It would be important to determine if the AID tag affects protein levels and localization in these cases. RNAi of *cul-1* or proteasome inhibition could test whether an endogenous ubiquitin ligase could interact with the AID tag in the absence of TIR1. Additionally, more information is required to determine rules for optimal AID tag placement in both structured and unstructured domains of proteins. As a precaution, we tend to use long 10-30 amino acid flexible linker sequences to space the AID* tag away from the protein of interest.

Intuitively, one might assume that TIR1 levels correlate with degradation efficiency. Indeed, evidence supports stronger depletion of AID*-tagged proteins when using TIR1 expressed from multi-copy arrays compared to single-copy transgenes (O. Hobert, personal communication). It will be important for groups to note expression levels of their AID fusion protein and the tissue-specific promoter when assessing efficiency of depletion. Mining a single-cell RNA-seq (scRNA-seq) datasets from L2 larvae (Cao *et al*. 2017), embryos (Tintori *et al*. 2016; Packer *et al*. 2019), and ideally, future stage-specific scRNA-seq datasets would allow identification of new, strongly expressed tissue-specific promoters. Other approaches to increasing the expression levels of tissue-specific promoters could be integrating arrays. Such an approach was required for optimal performance of the cGal system (Wang *et al*. 2017). A recent approach using CRISPR to allow site-specific integration of arrays could allow new multicopy TIR1 drivers to be inserted at specific chromosomal loci (Yoshina *et al*. 2016). Alternatively, multicopy tissue-specific Gal4 strains could be used to drive tissue-specific expression of UAS::TIR1, though this would increase the complexity of crosses to create new strains. However, we also note that effective depletion is possible even when TIR1 levels are low enough where the BFP reporter is undetectable (Figure 4). This result highlights the importance of functionally testing new TIR1 transgenes with FP::AID*-tagged alleles of interest.

The ability to rapidly deplete proteins with temporal and cellular resolution allows precise dissection of the roles of gene products in developmental processes of interest. With the ever-increasing efficiency of genome editing and continued refinement of the AID system, one can envision creating libraries of FP::AID*-tagged genes covering the genome and a bank of TIR1 strains to allow depletion in virtually all cell types.

## Acknowledgments

Many thanks to Matthew Schwartz for providing reagents and suggestions to get the SapTrap system established in our lab. We thank Oliver Hobert, Troy McDiarmid, and Francis McNally for helpful discussions about the AID system. We thank Alison Woollard for sharing unpublished information about a minimal SCM enhancer. G.A., M.T.L., H.N.S., B.D., J.M.R., and J.D.W. are supported by the NIH/NIGMS [R00GM-107345]. J.D.H. is supported by F32 GM131577, J.D.H. and B.G. are supported by R35 GM134838, and T.D. and D.J.R are supported by R01 GM121625. D.Q.M. is supported by the NIH NIGMS [1R01GM121597]. D.Q.M. is also a Damon Runyon-Rachleff Innovator supported (in part) by the Damon Runyon Cancer Research Foundation [DRR-47-17]. M.A.Q.M. is supported by the NIGMS [3R01GM121597-02S2]. N.J.P. is supported by the American Cancer Society [132969-PF-18-226-01-CSM]. T.N.M. is supported by the NICHD [F31HD100091]. Some strains were provided by the *Caenorhabditis* Genetics Center, which is funded by the NIH Office of Research Infrastructure Programs [P40 OD010440].

## AUTHOR CONTRIBUTIONS

D.J.D. designed the built-in F2A::BFP::AID*::NLS reporter strategy for TIR1 expression and activity. G.A, T.D., M.L., M.A.Q.M., J.D.H, N.J.P., D.Q.M., D.J.D., D.J.R., and J.D.W. conceived and designed the experiments. G.A. T.D., R.D., J.D.H., W.Z., T.N.M.-K., S.S.S., D.Q.M., D.J.D., D.J.R., and J.D.W. designed the constructs. T.D., J.D.H., R.D., N.J.P., J.M.R, and D.J.D. performed the microinjections. G.A., T.D., M.L., M.A.Q.M., J.D.H., N.J.P., and D.J.D. performed the crosses and characterized strains. G.A., T.D., M.L., R.D., M.A.Q.M., J.D.H., H.N.S., N.J.P., R.M., B.D., J.M.R., and D.J.D. performed the experiments. G.A., T.D., M.A.Q.M., J.D.H., N.J.P., D.Q.M., D.J.D., D.J.R., and J.D.W. analyzed and quantified the data. G.A. and J.D.W. wrote the manuscript with contributions from the other authors. The authors declare no competing interests.

## Figure Legends

**Figure S1.**
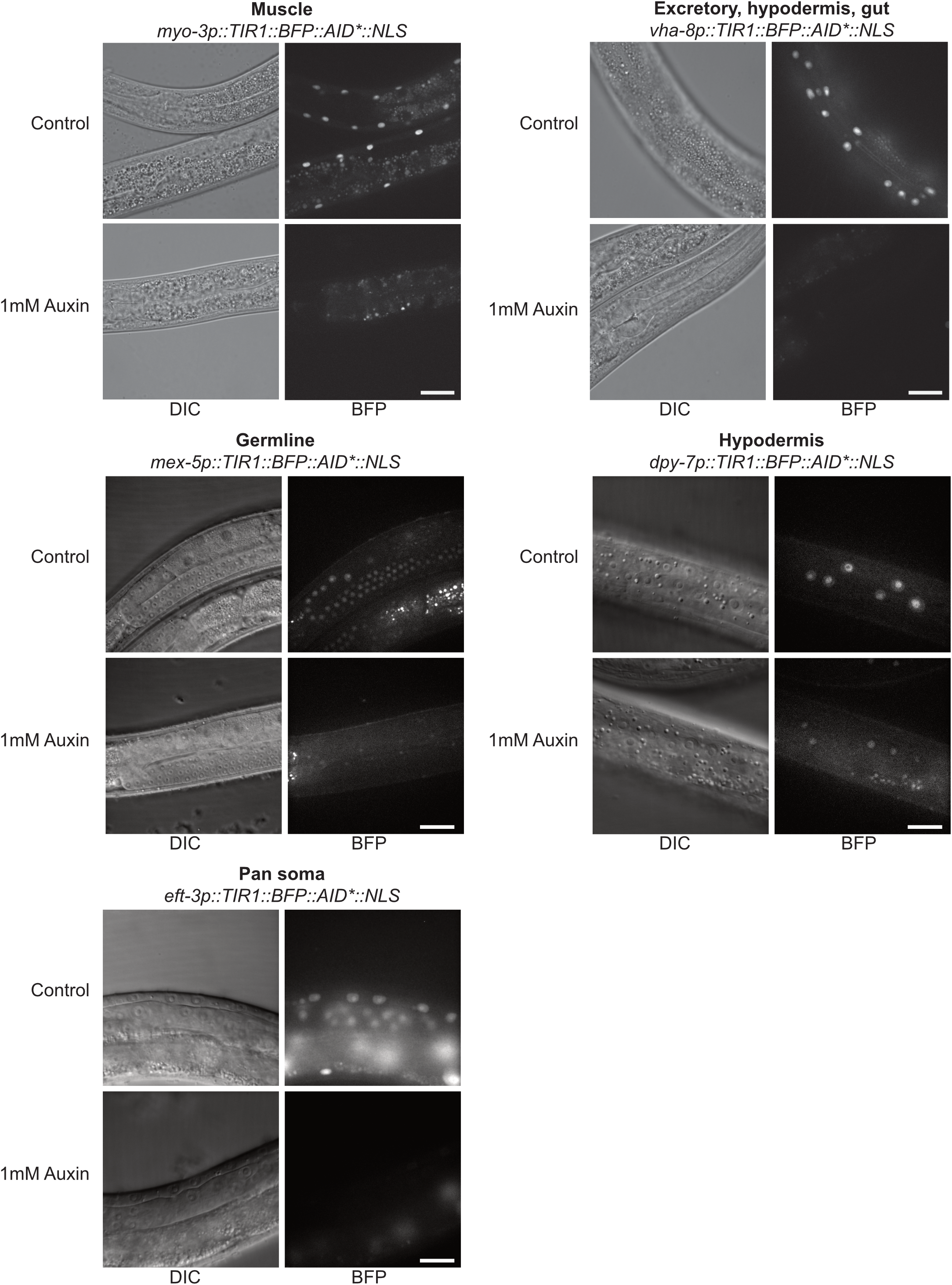
Functional test of new TIR1 expressing strains. Strains of the indicated genotypes were grown on 1 mM water-soluble synthetic auxin (potassium salt of 1-naphthaleneacetic acid; K-NAA) for ∼24 hours (*myo-3p* and *vha-8p)* or for one hour (*mex-5p, dpy-7p*, and *eft-3p*) before imaging. Animals were similarly grown on control seeded NGM plates lacking K-NAA. For all strains, the expected BFP expression pattern was observed in control animals, and BFP expression was reduced or lost in auxin treated animals. Scale bars represent 20 µm (*myo-3p, vha-8p*), 15 µm (dpy-7p, *eft-3p*), and 40 µm (*mex-5p*).

